# Mass spectrometry of RNA-binding proteins during liquid-liquid phase separation reveals distinct assembly mechanisms and droplet architectures

**DOI:** 10.1101/2022.09.28.509878

**Authors:** Cagla Sahin, Aikaterini Motso, Xinyu Gu, Hannes Feyrer, Dilraj Lama, Tina Arndt, Anna Rising, Genis Valentin Gese, Martin Hällberg, Erik. G. Marklund, Nicholas P. Schafer, Katja Petzold, Kaare Teilum, Peter G. Wolynes, Michael Landreh

## Abstract

Phase separation of heterogeneous ribonucleoproteins (hRNPs) drives the formation of membraneless organelles, but structural information about their assembled states is still lacking. Here, we address this challenge through a combination of protein engineering, native ion mobility-mass spectrometry, and molecular dynamics simulations. We used a phase separation-compatible spider silk domain and pH changes to control the self-assembly of the hRNPs FUS, TDP-43, and hCPEB3, which are implicated in neurodegeneration, cancer, and memory storage. By releasing the proteins inside the mass spectrometer from their native assemblies, we could monitor conformational changes associated with phase separation. We find that NT*-FUS monomers undergo an unfolded-to-globular transition, whereas NT*-TDP-43 oligomerizes into partially disordered dimers and trimers. NT*-hCPEB3, on the other hand, remains fully disordered with a preference for fibrillar aggregation over phase separation. The divergent assembly mechanisms result in structurally distinct complexes, indicating differences in RNA processing and translation depending on biological context.

## Introduction

Liquid-liquid phase separation (LLPS) of proteins into liquid droplets is the governing principle for the formation of membraneless organelles and controls diverse biological processes from ribosome assembly to RNA processing ^1,2^. Genome-wide analyses have revealed that the ability to form liquid droplets via LLPS is a common feature of the proteome ^3^. These findings raise the interesting possibility that LLPS is a generic property of polypeptide chains ^4^, as suggested previously for the capacity to form amyloid-like fibrils ^5^. Interestingly, both LLPS and amyloid formation have been observed for heterogeneous ribonucleoproteins (hRNPs). hRNPs are a diverse family of proteins with a modular architecture composed of folded RNA-recognition motifs (RRMs) and disordered low-complexity domains (LCDs) with prion-like properties ^6,7^. hRNPs readily form liquid droplets *in vitro* and *in vivo*, and the biological relevance of their droplet states for the cellular RNA metabolism have been clearly established ^8^. The perhaps best-understood hRNPs are Fused in Sarcoma (FUS) and Transactive Response DNA-binding protein 43 (TDP-43), which are components of stress granules and involved in the cellular RNA metabolism. Both proteins have a strong propensity to form fibrillar aggregates *in vitro* and *in vivo*^9–11^ and have been identified as major components in neuronal inclusion bodies from patients with amyotrophic lateral sclerosis (ALS), and, for TDP-43, also in patients with limbic-age related TDP-43 encephalopathy (LATE), which has symptoms similar to Alzheimer’s Disease ^8,12,13^. It has been proposed that aberrant LLPS can give rise to toxic protein aggregates and thus become a driving factor of neurodegenerative processes ^6^. On the other hand, fibril formation has been identified as a functional feature of the human cytoplasmic polyadenylation element binding protein 3 (hCPEB3), an hRNP that is a key regulator of synaptic plasticity and long-term memory formation ^14^. Its homologues from *Drosophila melanogaster* (Orb2), *Aplysia californica* (apCPEB), and mouse (mCPEB3) are functional prions that regulate transcriptional activity by assembling into fibril-like structures ^14^. Yet several CPEB proteins have been found to also undergo LLPS, including mCPEB3, hCPEB3, and Orb2, albeit only under specific conditions such as SUMOylation ^15^, in crowding agents ^16^, or as precursor of fibril formation ^17^.

The fact that several hRNPs can adopt multiple competing assembly states raises the question whether there are distinct conformational signatures that can be associated with LLPS. Identifying structural features that distinguish pathogenic from physiological assemblies would be an important step towards pharmacological intervention in toxic aggregation. Although the sequence requirements for LLPS are comparatively well studied ^2,4,18^, we lack an understanding of how these translate into a three-dimensional architecture. NMR spectroscopy has provided valuable information ^19^, however, obtaining and comparing structural determinants of droplet formation remains challenging due to the poor solubility and low conformational stability of many hRNPs. For example, full-length FUS and TDP-43 require the presence of strong denaturants or fusion to expression tags which prevent aggregation as well as LLPS, rendering purified proteins either non-native or non-functional ^20–23^. Furthermore, the choice of renaturation strategy can affect the balance between LLPS and aggregation, leading to conflicting observations ^24,25^.

Here we develop a pH-responsive LLPS system by fusing aggregation-prone hRNPs to an engineered spider silk domain and use native mass spectrometry (nMS) in combination with ion mobility spectroscopy (IM) to identify conformational changes that are associated with droplet formation. Using MD simulations, we find that despite similar domain organizations, hRNPs adopt distinct conformational states during assembly, which affect the orientation of bound RNAs and may distinguish specific biological contexts.

## Results

### A spider silk tag enhances hRNP stability without abolishing LLPS

We asked how LLPS of severely aggregation-prone proteins is controlled in nature. Major ampullate spidroins (MaSp), the proteins that make up spider dragline silk, undergo amyloid-like aggregation during spinning, yet are stored at high concentrations in the sac of the silk gland ^26^. The stored proteins display hallmarks of phase separation *in vivo* and contain LC domains that mediate LLPS *in vitro*^27^. We previously designed an artificial mini-spidroin (miniMaSp1) composed of a regulatory N-terminal (NT) domain, which cross-links the fibers at low pH, a C-terminal domain which acts as aggregation switch, and two LC repeats (Figure 1a)^28^. Brightfield microscopy reveals that at pH 7.5, miniMaSp1 forms spherical droplets with a diameter of 4-8 μm, in agreement with previous reports of LLPS by mini-spidroins ^27^. The NT domain, albeit highly soluble on its own ^29^, does not appear to inhibit droplet formation. Interestingly, the amino acid compositions of the LC regions of FUS and TDP-43 closely resemble that of MaSp1, except for poly-Ala blocks that are specific for spidroins (Figure 1b). We therefore speculated whether fusion to a charge-reversed NT mutant (D40K/K65D), termed NT*, which does not cross-link at low pH ^30^, could enable the production of soluble full-length hRNPs without interfering with LLPS *in vitro* (Figure 1c). Indeed, recombinant expression in *E. coli* yielded high amounts of predominantly soluble NT*-FUS and NT*-TDP-43 (Figure S1a). We observed loss of soluble NT*-TDP-43 after incubation in common buffers, but good solubility in water, in line with previous findings (Figure S1b) ^31^. We screened pH stability by incubating the proteins in H_2_O at pH values ranging from 6 to 11 and monitored the formation of insoluble aggregates by centrifugation and SDS PAGE (Figure 1d). We found that the fusion proteins are stable in H_2_O / ammonia or H_2_O / imidazole at pH regimes above their pI (Figure 1d, e). We then asked whether the purified proteins retain natively folded RNA-binding domains. Denaturing polyacrylamide gel electrophoresis (PAGE) of NT*-FUS revealed a large amount of co-purified nucleic acids which were sensitive to RNase A treatment by reduction of overall intensity of the band, indicating that the protein retains bound RNAs from the expression host (Figure S1c), confirming that the RNA binding capacity was at least partial intact. In the following, residual RNA was reduced by a 1 M NaCl washing step. To test the ability of the fusion proteins to undergo LLPS, we diluted NT*-FUS from a pH 9 dH_2_O stock into 20 mM Tris pH 6, 500 mM NaCl, conditions under which full-length FUS undergoes phase separation ^32^ and monitored it with microscopy. We clearly observed spherical droplets that fuse on a minute timescale (Movie S1), which demonstrates that LLPS of NT*-FUS purified under native conditions can be triggered by adjusting buffer conditions and is not inhibited by the NT*-tag.

**Figure 1.**
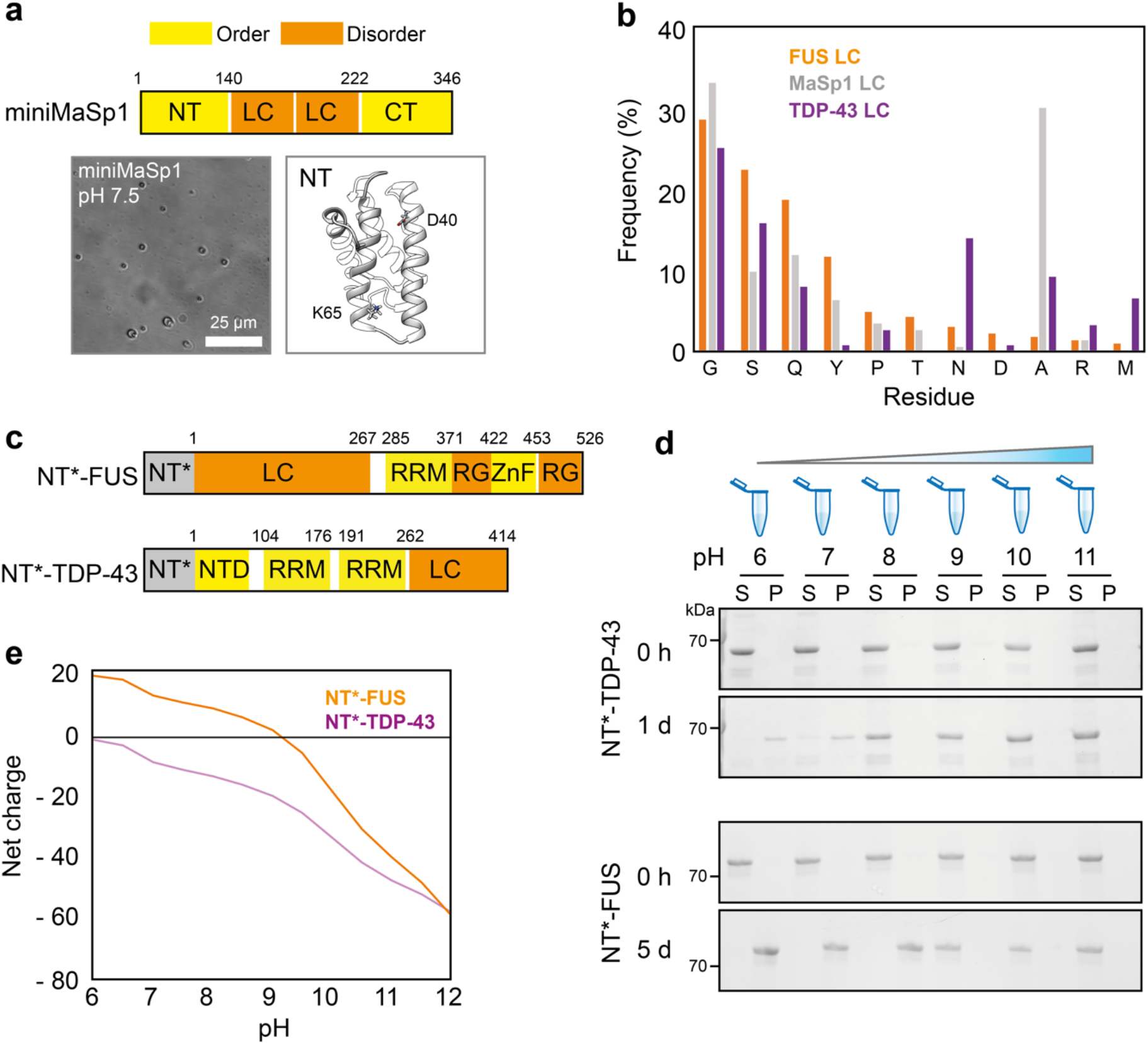
Design and characterization of chimeric hRNPs. (a) The architecture of the miniMaSp1 spidroin with NTD, N-terminal domain; LC, low complexity region, and CTD, C-terminal domain. Brightfield microscopy shows that miniMaSp1 forms spherical droplets at pH 7.5. The structure of the NT domain (PDB ID 4FBS)^33^ is rendered as cartoon with the residues K65 and D40, which are swapped in the NT* variant, shown as sticks. (b) The LC domains from FUS, TDP-43, and MaSp1 have similar amino acid compositions, except for poly-A blocks that are specific for MaSp proteins. (c) Architecture of the NT*-tagged full-length FUS and TDP-43 variants used in this study. RRM, RNA recognition motif; ZnF, zinc finger. (d) NT*-FUS and NT*-TDP-43 resist aggregation and display long-term stability under mildly alkaline conditions. Solubility was determined by incubating protein at pH values between 6 and 11 and separating soluble (S) and insoluble (P) fractions by centrifugation. (e) Computed net charge and solubility transitions of NT*-FUS and NT*-TDP-43 shows aggregation close to the respective pIs (8.93 for NT*FUS and 6.05 for NT*TDP-43). The net charges were predicted using the Pro pKa server (http://propka.ki.ku.dk/pka/2khm.html).

### NT*-FUS and NT*-TDP-43 in liquid droplets can be observed by nMS

Having established production of full-length NT*-FUS and NT*-TDP-43 under native conditions, we asked whether we could observe specific structural changes associated with LLPS. For this purpose, we turned to nMS. In nMS, intact protein complexes are ionized through electrospray ionization (ESI) and gently transferred to the gas phase for mass measurements without disturbing non-covalent interactions ^34^. The number of charges that is acquired during ESI correlates with the surface area and flexibility of the protein in solution ^35,36^. By combining nMS with IM, we can determine the collision cross-section (CCS) of the ionized proteins, and in this manner obtain insights into the conformational preferences and relative stabilities of disordered proteins in the gas phase, and how they are related to solution structures ^35,37^. We have previously used nMS to reveal soluble intermediates in spider silk formation ^38^. Like silk assembly, LLPS can be induced in the absence of salt ^32^ and controlled by adjusting the solution pH ^22^. As LLPS and nMS additionally require a similar protein concentration in the low micromolar range, we devised a two-pronged approach in which we prepared NT*-tagged hRNPs at different pH regimes, monitored LLPS formation by brightfield microscopy, and subjected the same samples to nMS analysis (Figure 2a). As solvent system, we chose water/ammonia, which has been shown to be suitable for preparation of the aggregation-prone TDP-43 ^31^. Importantly, the same system is well suited to study protein folding by nMS. Unlike acidic conditions, alkaline pH does not induce additional unfolding during ESI, allowing for an accurate assessment of a protein’s folded states in response to pH ^39^.

**Figure 2.**
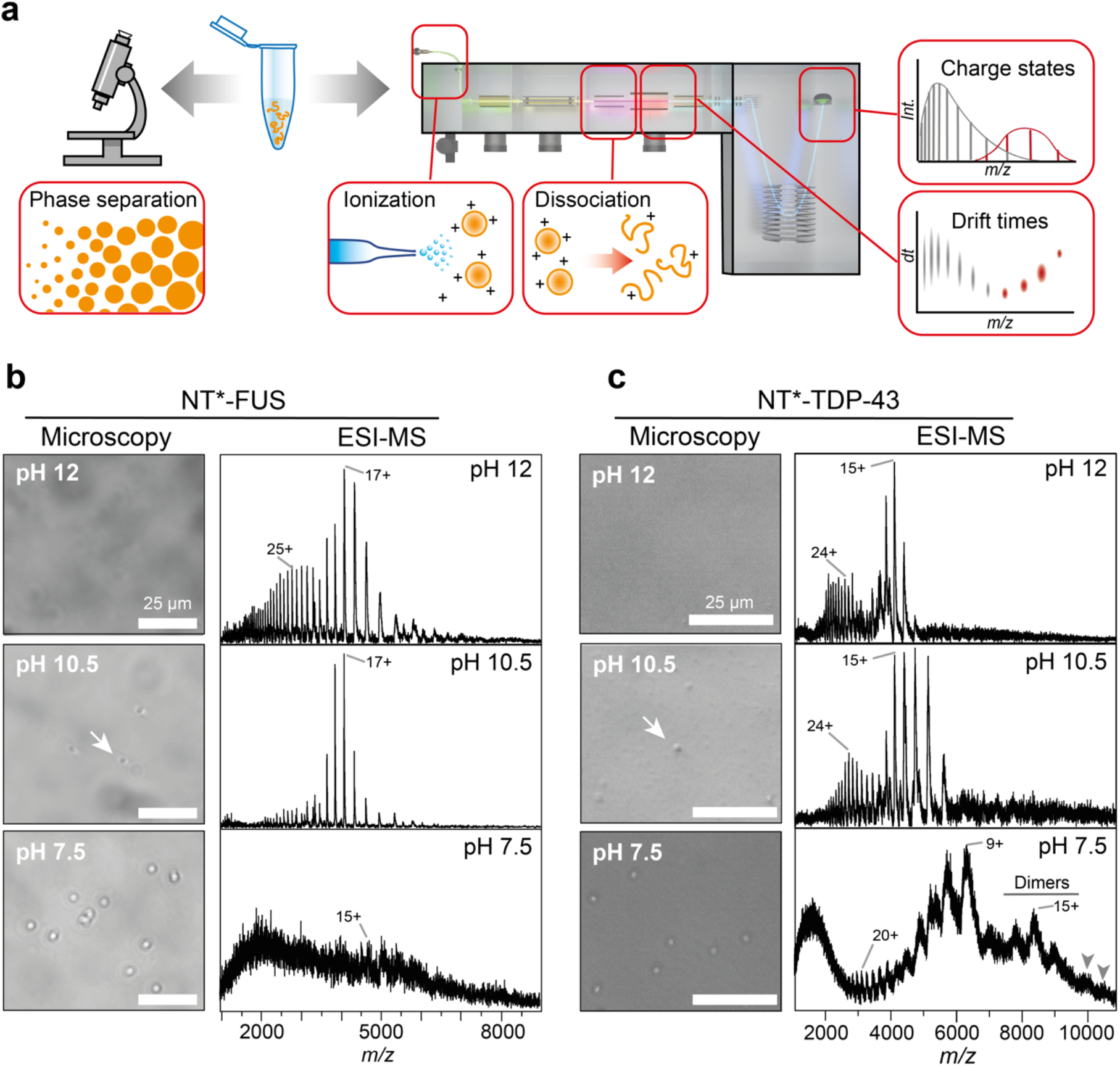
Microscopy and MS of NT*-FUS and NT*-TDP-43 under denaturing and LLPS conditions. **(a)** Summary of the combined microscopy and nMS approach. NT*-tagged hRNPs are exposed to different pH regimes under salt-free solution conditions and subjected to both brightfield microscopy and MS analysis. By combining nMS with IM and gas-phase dissociation, it is possible to extract ion charge states and drift times for soluble and assembled hRNPs. **(b)** Brightfield microscopy of NT*-FUS shows the onset of droplet formation at pH 10.5 (arrow) and from nMS it appears as complete LLPS at pH 7.5. nMS reveals a shift from a broad, bimodal to a narrow, monomodal CSD between pH 12 to 10. At pH 7.5, only traces of monomeric NT*-FUS can be detected by nMS. **(c)** NT*-TDP-43 forms a few vaguely defined droplets at pH 10.5 (arrow) and undergoes complete LLPS at pH 7.5. nMS shows a bimodal CSD at pH 12 and 10.5. At pH 7.5, a pronounced shift to lower charges occurs, and peaks corresponding in mass to dimers and trimers (arrows indicate the 19+ and 18+ ions of the NT*-TDP-43 trimer) can be detected.

To test whether we can detect proteins in liquid droplets by MS, we chose NT*-FUS, since FUS is often used as a model system for LLPS. Starting at pH 12 (Figure 2b), microscopy showed no discernible structures, as expected from a visible soluble fraction after 5d at pH >9 (Fig. 1d). nMS analysis revealed well-resolved peaks with a bimodal charge state distribution (CSD) centered on the 25+ and 17+ ions. Broad CSDs are indicative of conformationally flexible proteins, where extended states give rise to highly charged ions, while compact states preferentially acquire low charges.^37^ At pH 10.5, we observed by microscopy very few spherical assemblies with poor contrast, possibly representing the onset of droplet formation. Strikingly, the mass spectrum showed a pronounced shift towards a monomodal CSD around the 17+ ion with a decrease in pH, while the higher-charged distribution disappeared. At pH 7.5, we detected well-resolved droplets with a diameter of around 5 μm (Figure 2b) that are morphologically indistinguishable from those formed in 20 mM Tris, 500 mM NaCl, pH 6 (Movie S1). This shift was accompanied by a near-complete loss of protein signal in the mass spectrum, in line with the incorporation of virtually all NT*-FUS molecules into assemblies that are not detectable by MS. To test this possibility, we employed collisional activation in the ion trap of the mass spectrometer. Collision-induced dissociation breaks down protein assemblies that are too large or heterogeneous for detection by MS, releasing individual components which can be readily observed (Figure 2a) ^40^. Indeed, activation resulted in the appearance of peaks that correspond to monomers of the intact NT*-FUS protein (Figure S2). This observation supports our hypothesis that NT*-FUS self-assembles without adopting a specific stoichiometry.

Next, we examined NT*-TDP-43 (Figure 2c). As for NT*-FUS, no droplets could be observed by microscopy at pH 12. nMS showed a bimodal CSD with a compact charge state envelope around the 15+ ion, as well as highly charged ions with lower intensity, confirming the presence of flexible, monomeric protein. At pH 10.5, we observed sparse spherical assemblies by microscopy that resemble those seen for NT*-FUS. However, the bimodal CSD in nMS remained largely unchanged compared to pH 12. Upon lowering the pH to 7.5, we find NT*-TDP-43 formed well-defined droplets. Unlike NT*-FUS, however, the protein could still be detected by nMS without collisional activation. Although the CSD remained broader than for NT*-FUS, it shifted towards the higher m/z region. We furthermore observed peaks corresponding in mass to NT*-TDP-43 dimers, as well as traces of trimers (Figure 2c, S3). We conclude that in contrast to NT*-FUS, NT*-TDP-43 assembles into dimers, trimers, and possibly higher oligomers, under LLPS conditions.

### Distinct conformational signatures of NT*-FUS and NT*-TDP-43 during LLPS

We asked if the different CSDs of NT*-FUS and NT*-TDP-43 are caused by conformational changes during LLPS. Previous studies have shown that the flexibility of disordered proteins in solution is reflected in the balance of Coulombic stretching and collapse that the proteins experience in the gas phase ^35^. By comparing the distributions of CCSs and charge states, we can therefore determine whether a protein is more likely to be compact (narrow CSD with near-constant CCS) or extended (broad CSD and wide CCS range) in solution. We therefore used IM-MS to determine CCSs of NT*-FUS and NT-TDP-43 at each pH value. First, we measured the CCS of NT*-FUS at pH 12. Plotting the CCS as a function of ion charge revealed a steep rise in CCS as the charge state of the protein increases (Figures 3a). This finding suggests that the CCS of the protein at alkaline pH is determined mainly by its charge, as expected for a fully disordered protein.^35^ At pH 10.5, where we observed a narrow, monomodal CSD around 17+, the CCSs remains nearly constant for all major charge states. The most abundant charge states of 16+ and 17+ display similar CCSs of 4676 and 4606 Å^2^, respectively, and considerably narrower arrival time distributions (Figure 3b). Taken together, the changes in CSD and CCS closely resemble the behavior of globular proteins at this pH ^39^, which leads us to conclude that NT*-FUS transitions from a flexible, disordered state to a compact conformation as the pH is lowered. The IM-MS data are thus in agreement with recent NMR and EPR studies showing that FUS undergoes a significant compaction as it approaches LLPS ^41^.

**Figure 3.**
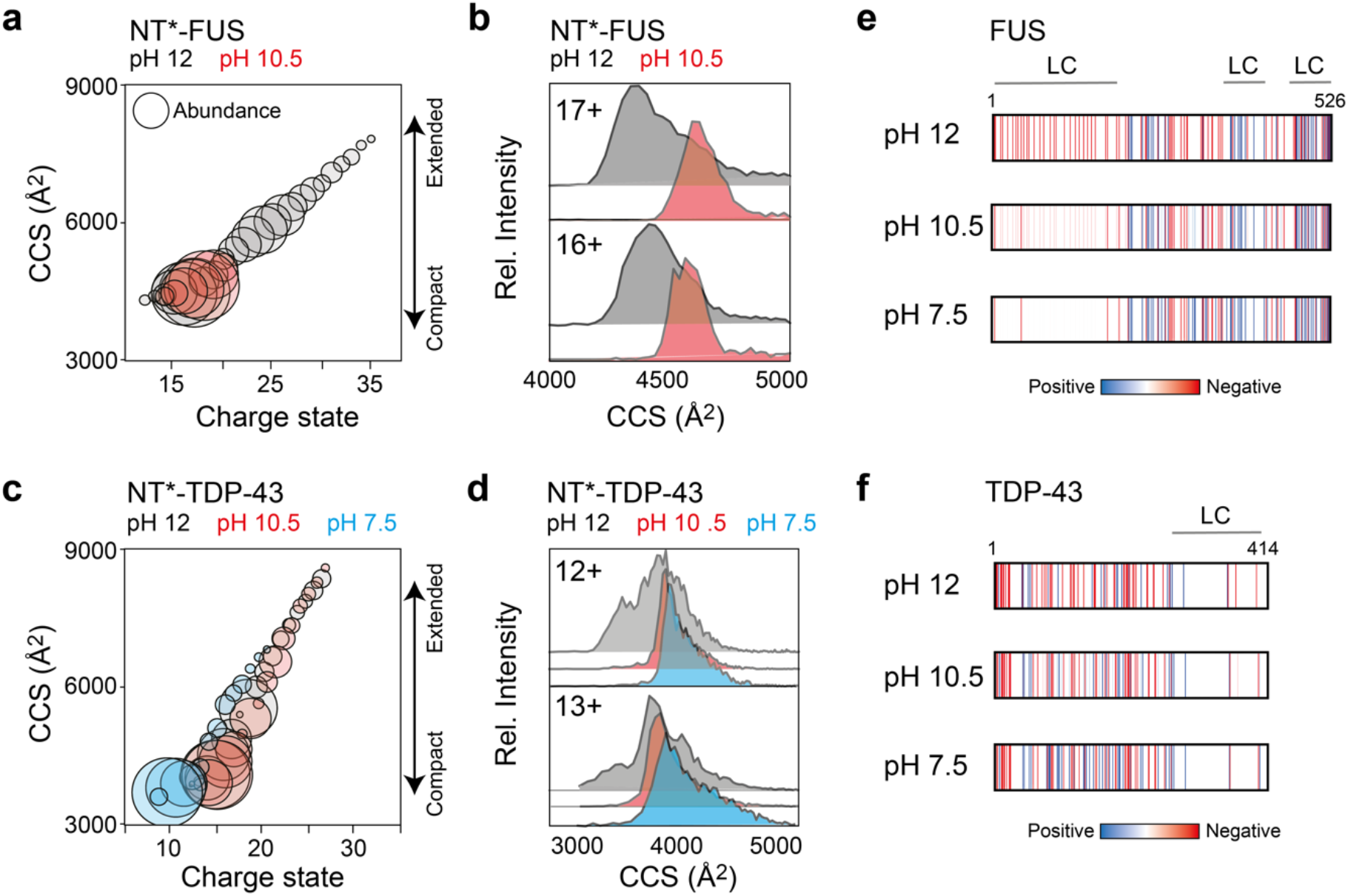
Structural changes during LLPS of NT*-FUS and NT*-TDP-43. **(a)** Plotting the CCS of NT*-FUS as a function of charge state shows a steep rise in CCS with increasing ion charge at pH 12 (grey). At pH 10 (red), all ions display similar CCS values which are less dependent on ion charge, which is a hallmark of compact proteins. Ion abundances are indicated by the diameter of each circle. **(b)** CCS distributions for the 17+ and 16+ charge states of NT*-FUS are narrower at pH 10.5 compared to pH 12. The slight increase in CCS may indicate that NT*-FUS adopts a more defined structure at pH 10.5. **(c)** The charge state-CCS plot for NT*-TDP-43 shows a similar dependence of CCS on ion charge for pH 12, 10.5, and 7.5, indicating no pronounced unfolded-to-globular transition. **(d)** CCS distributions for the 12+ and 13+ ions of NT*-TDP-43, the charge states that can be detected at all pH values, show no pronounced changes in peak width or CCS at any condition. **(e, f)** Plotting the distribution of charged residues (based on their pKa) in the sequences of FUS and TDP-43 at pH 12, 10.5, and 7.5 shows that the N-terminal LC domain of FUS shifts from negative to neutral charge as the pH is lowered. TDP-43, on the other hand, does not display notable shifts in charge distributions.

We then used the same strategy to analyze NT*-TDP-43 at pH 12, 10.5, and 7.5. At pH 12 and 10.5, the CCS-charge state plot reveals that the protein’s cross section increases with ion charge in a similar manner as NT*-FUS at pH 12 (Figures 3c). At pH 7.5, we detected lower charge states and CCSs, but also a population with high charges and CCSs. Furthermore, the 12+ and 13+ ions, the lowest charge states present in all conditions, had similarly low CCSs, with only moderately narrower arrival time distributions at low pH (Figure 3d). From the CSD and CCS data, we conclude that NT*-TDP-43 does not undergo a pronounced unfolded-to-globular transition like NT*-FUS. Instead, its response to lowered pH is consistent with the behavior of a partially disordered protein ^39^. Interestingly, we also detected higher oligomeric states for NT*-TDP-43 at pH 7.5 that were absent in NT*-FUS. To obtain more structural information on oligomerization, we further examined the dimer by IM-MS. The CCS of the dimer was found to be 6812, 7009, and 7143 Å^2^ for the 14+, 15+, and 16+ charge states, respectively. By combining CCS values with oligomeric state and molecular weight information to mine the PDB, it is possible to extract likely complex shapes ^42^. However, this strategy did not yield a clear preference, but rather suggests a range of oblate or prolate shapes (Figure S3). Taken together, we conclude that monomers and oligomers of NT*-TDP-43 retain some conformational heterogeneity.

The IM-MS data suggest that NT*-FUS and NT*-TDP-43 display distinct structural features in IM-MS as they approach the LLPS regime: NT*-FUS undergoes significant compaction around pH 10.5 and is subsequently completely incorporated into LLPS assemblies as the pH is lowered to 7.5. NT*-TDP-43, on the other hand, remains flexible, but shows stepwise oligomerization at pH 7.5. Barran and co-workers have reported that the conformations of disordered proteins in nMS are largely governed by charge pattern ^43^. We therefore computed the distribution of charged residues along the FUS and TDP-43 sequences at each pH (Figure 3c). We find that the LC domain of FUS, which is rich in tyrosine, undergoes a shift from negative to neutral charge as the pH drops below the pKa of tyrosine at 10.4. For TDP-43, on the other hand, we do not observe pronounced changes in charge pattern, since both positively and negatively charged residues are distributed relatively evenly throughout the sequence.

### Structural models of FUS and TDP-43 assembly intermediates

To understand how the specific structural preferences of FUS and TDP-43 mediate self-assembly, we devised a hybrid strategy combining AlphaFold2 (AF2) structure prediction and molecular dynamics (MD) simulations (Figure S4). For FUS, we used AF2 to obtain full-length models of monomeric FUS, which we subjected to all-atom MD simulations in solution at pH 7.5. Starting from an extended, random conformation generated by AF2, FUS adopts a partially compact structure with intact RRM during a 500 ns simulation in solution (Figure 4a). Strikingly, the tyrosine-rich N-terminal LCD and the glycine/arginine-rich C-terminal LCDs remain mostly segregated in the model. The resulting structure shows distinct lateral distribution of arginines and tyrosines, with a Tyr- and an Arg-rich pole (Figure 4a). Most tyrosines and arginines are located at the surface of the protein. However, we also found intramolecular contacts at the intersection near the RRM. We detected contacts between Tyr 14, 38, 148, and 177 with Arg 244, 218, 213, and 248, respectively (Figure 4a). These interactions likely contribute to the compaction of the protein in solution by connecting the N- and C-terminal LCDs, with the RRM sandwiched in between.

**Figure 4.**
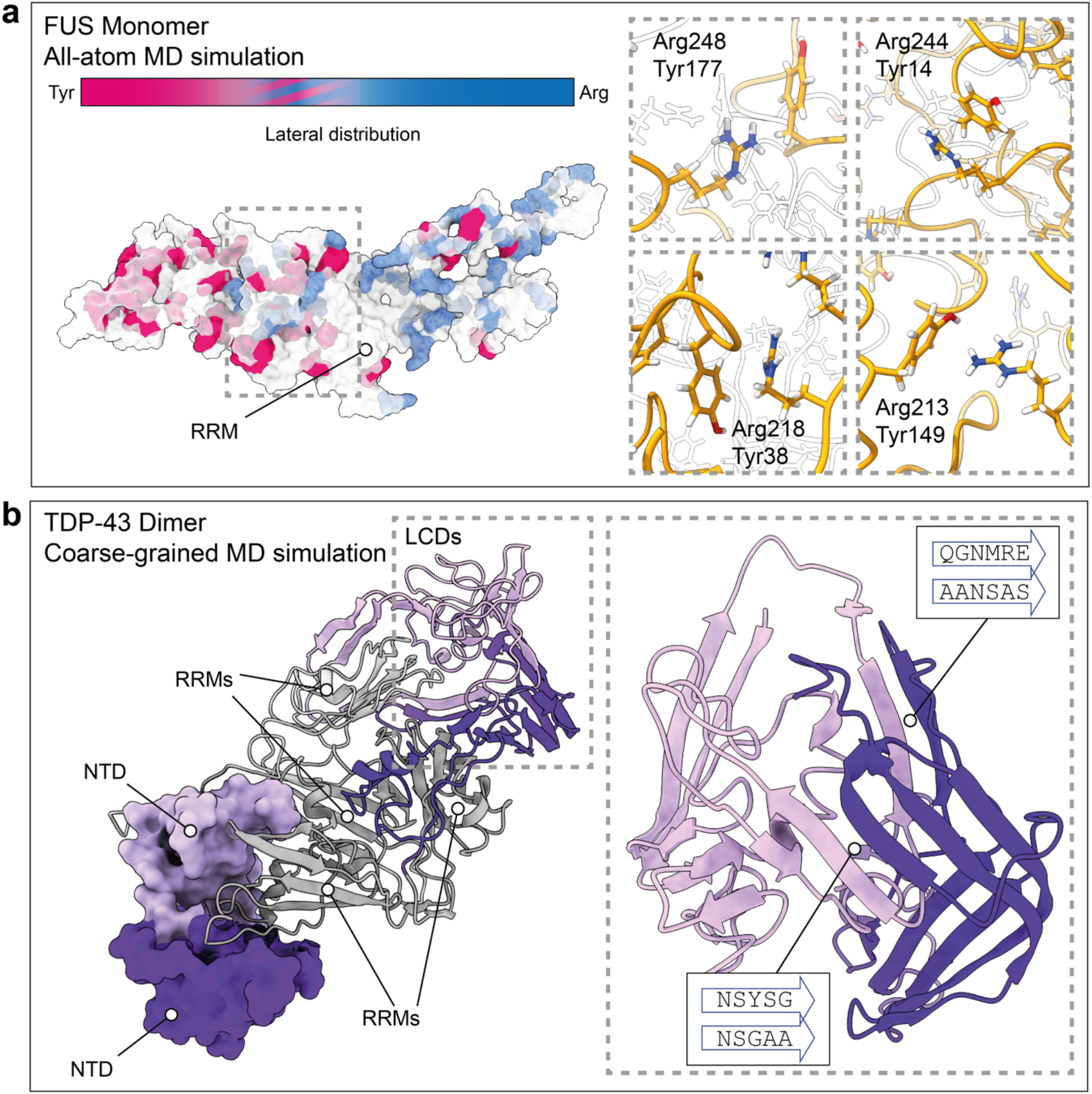
Models of FUS and TDP-43 species observed by MS. **(a)** All-atom MD of monomeric FUS shows a compact conformation and a bipartite organization with a tyrosine- and an arginine-rich domain. While most arginine and tyrosine residues are located at the surface, we also detect contacts between N- and C-terminal LCDs mediated by Arg-Tyr interactions, indicated by a dashed box. **(b)** A representative example of the AF2- and coarse-grained MD-derived model of dimeric TDP-43. The end structure from two rounds of cooling shows tight interactions between the folded NTDs (shown as surface representations). The RRMs (grey) are in proximity mediated by intermolecular contacts between the LCDs. Insert: Representative LCD interactions between two protomers reveal extensive inter- and intramolecular β-sheet formation. Sequences engaged in the intermolecular interactions are indicated in boxes.

Next, we developed models of monomeric, dimeric, and trimeric TDP-43 using coarse-grained MD simulations in solution at pH 7. Since TDP-43 is known to oligomerize via its N-terminal domains ^44,45^, we used AF2 to build NTD dimers and trimers, which we used to align the NTDs of monomeric TDP-43 models derived from SAXS measurements ^31^. To extract structural preferences, we cooled the models of fulllength TDP-43 monomers, dimers, and trimers from 400 to 300 K with 20 replicates, under AWSEM force field ^46^. These 1^st^ generation models were subjected to a second round of cooling and the resulting 2^nd^ generation models were used to compute contact maps and electrostatic energies (Figure S4). Comparison of the 20 end structures reveals a significant degree of compaction but no convergence to a single conformation. The contact maps show that the RRMs interact with the NTDs and LCDs, but there are almost no contacts between NTDs and LCDs (Figure S4). The fold and interactions of the NTDs are preserved in all models (Figure 4b, Figure S4), suggesting that the NTDs can mediate oligomerization of full-length TDP-43. Strikingly, the C-terminal LCDs consistently fold into β-sheet-rich structures that bring the RRMs in dimers and trimers into proximity (Figure 4b, Figure S4). The C-terminal segments involved in β-sheet formation vary between replicates but always include prion-like sequences with high aggregation propensity (Figure 4b). Interestingly, in the trimer, we find that two neighboring LCDs interact whereas the third LCD points, which is probably a result of the curvature of the NTD trimer (Figure S4). The orientation of RRMs and LCDs varies between replicates, which indicates significant flexibility, in agreement with the IM-MS results (Figure S4). Lastly, we calculated the electrostatic energies of the end structures, and found them to be slightly more favorable in the oligomers at neutral pH (Figure S4). This difference suggests that NTD interactions, which are charge-based ^45^, are favorable for TDP-43. Interactions between LCDs, on the other hand, occur almost exclusively in chargeless regions and are therefore pH insensitive. Taken together, the models reveal that TDP-43 forms flexible oligomers mediated by well-defined protein-protein interactions between NTDs and low-specificity contacts between LCDs.

### The conformational balance of hCPEB3 favors aggregation over LLPS

Encouraged by these results, we turned to a less-well understood hRNP, and asked whether nMS could reveal the structural preferences of native hCPEB3. As for FUS and TDP-43, tagging hCPEB3 (isoform 1) with NT* (Figure 5a) resulted in increased solubility, allowing us to purify the fusion protein under native conditions (Figure S5). However, NT*-hCPEB3 still displayed low stability, becoming immediately insoluble if the pH was lowered below 8 (Figure 5b). To better understand how NT* affects hCPEB3 aggregation, we conducted all-atom MD simulations of NT* fused to the first 40 residues of hCPEB3 (Figure S5). We observed transient contacts between the NT* surface and the polyglutamine stretch between residues 10-26, leading us to speculate that the NT* domain may reduce, but not block, self-association of this region, and thus retard hCPEB3 aggregation. Next, we examined LLPS of NT*-hCPEB3 with brightfield microscopy under the same conditions as for NT*-TDP43 and NT*-FUS. Strikingly, we did not see spherical droplets at lower pH, but instead elongated aggregates, which stained positive for Thioflavin T (ThT), a marker for amyloid structures (Figure S5), as reported for the *Drosophila* and *Aplysia* homologues.^47,48^ In line with the fluorescence microscopy, transmission electron microscopy (TEM) confirmed the presence of fibrillar aggregates (Figure 5b). nMS analysis of NT*-hCPEB3 showed a large population of highly charged ions, suggesting mostly extended protein conformations. At lower pH, the highly charged population remained dominant, whereas the signal/noise ratio in the spectra decreased significantly (Figure 5c). Plotting the charge distribution revealed that the N-terminal LC domain of hCPEB3, which has been identified as an aggregation hotspot, is virtually unaffected by changes in pH due to the near-complete absence of titratable residues (Figure 5d). In fact, the observations for NT*-hCPEB3 closely mirror the behavior of amyloidogenic peptides in nMS ^49^. Our results thus indicate that despite having a similar architecture as FUS and TDP-43, hCPEB3 favors aggregation over LLPS. The fact that the protein has a high aggregation propensity but can undergo conditional phase separation could indicate competing structural preferences, in line with the reported shifts in assembly state that hCPEB homologues can undergo *in vivo* ^15,48,50^. Importantly, we carefully considered the potential impact of the NT* tag on LLPS and aggregation and conclude that the observed structural plasticities are specific for each hRNP and unlikely to be affected by the presence of the NT* domain (see supplementary discussion).

**Figure 5.**
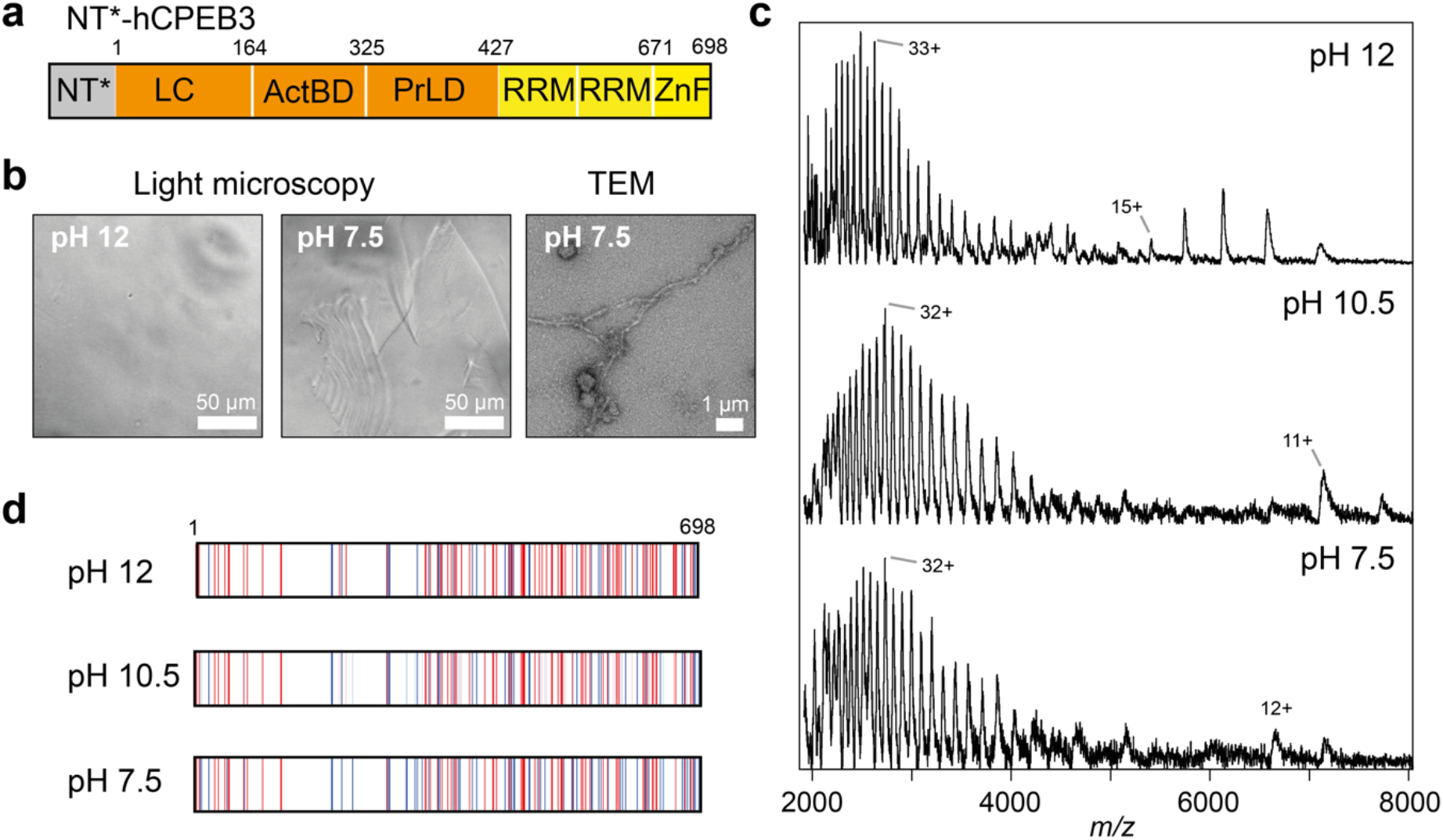
NT*-hCPEB3 remains disordered and undergoes aggregation at pH 8. **(a)** The architecture of NT*-hCPEB3. LC, low-complexity region; RRM, RNA recognition motif; ZnF, zinc finger; ActBD, actin-binding domain; PrLD, prion-like domain. **(b)** Brightfield microscopy of NT*-hCPEB3 at pH 12 and 7.5 shows the appearance of elongated aggregates at low pH. TEM shows fibrillar as well as some spherical aggregates. **(c)** nMS reveals highly charged monomeric NT*-hCPEB3 across the pH range tested. Note the decreased signal/noise ratio as the pH is decreased. **(d)** The predicted distribution of charged residues in the hCPEB3 sequence shows no pronounced local charges in response to pH.

## Discussion

Multiple hRNPs with similar domain organization undergo LLPS *in vitro* and *in vivo*,but insights into their assembled structures remain scarce. Here, we use the NT* domain to produce the human neuronal proteins FUS, TDP-43, and hCPEB3 under non-denaturing conditions. Native IM-MS shows that all three proteins populate different conformational states when LLPS is induced by lowering the pH. Using the insights from MS to inform MD simulations, we can delineate distinct assembly mechanisms (Figure 6). Importantly, these differences persist in presence of the NT* domain in all constructs, indicating that they are specific for each hRNP (see supplementary text).

**Figure 6.**
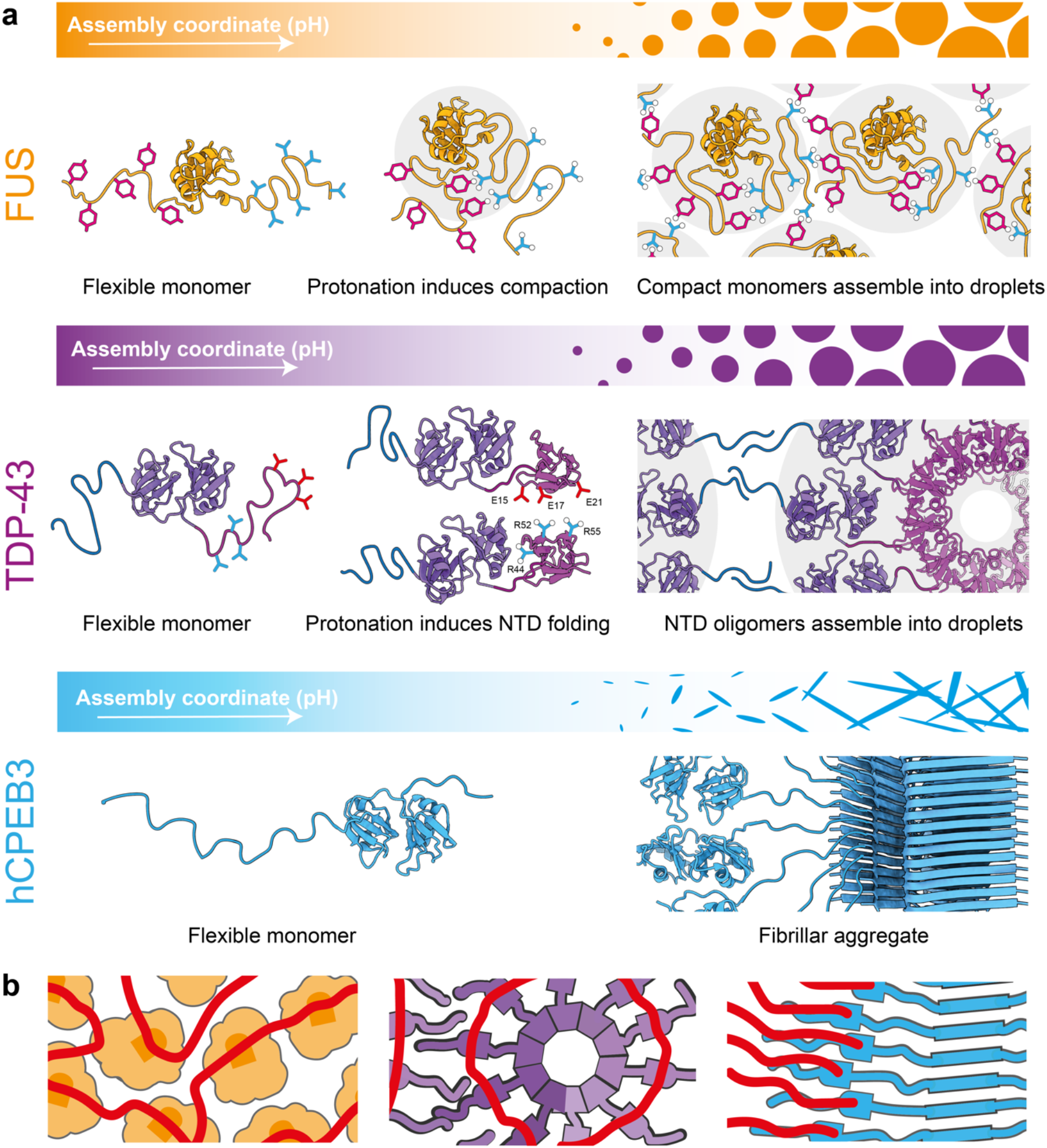
Divergent architectures and possible RNA binding modes of FUS, TDP-43 and hCPEB3 assemblies. (a) pH-dependent assembly of FUS is accompanied by protonation of tyrosine residues and increased intramolecular interactions that lead to compaction. The compact FUS monomers can then assemble into droplets via low-specificity contacts between surface-exposed arginine and tyrosine residues. TDP-43 contains an N-terminal domain which folds at pH 10 and subsequently oligomerizes via charge interactions. Oligomerization brings the C-terminal LC domains, which do not contain titratable residues, into proximity, and thus enables the interactions that drive LLPS. hCPEB3 remains mostly disordered across the pH range tested, but forms fibrillar aggregates. (b) Presumed orientation of RNAs in FUS, TDP-43, and hCPEB3 assemblies. RNA (red) binds the RRMs in compact FUS and this aligns the protein. TDP-43 adopts a helical structure, in which neighboring RRMs bind RNA target sequences. hCPEB3 aligns its RRMs along the fibril axis which serve as anchor points for RNA molecules

FUS has a bipartite organization, with a tyrosine- and an arginine-rich domain at the N- and C-termini, respectively. Cation-πinteractions between the tyrosine and arginine residues result in low-specificity interactions that drive LLPS ^18,51^. Importantly, these interactions are affected by the protonation state of tyrosine and can be disrupted at high pH ^22,52^. We now find that at high pH, FUS appears disordered, but adopts a compact state as the pH approaches the physiological range. At pH 7.5, the protein is completely incorporated in liquid droplets, which can be easily dissociated to release FUS monomers. MD simulations show that compaction of the protein results in a polar organization with an arginine- and a tyrosine-rich side. Together, these findings suggest that lowering the pH promotes cation-πinteractions between the C- and N-terminal domains, which give rise to a compact, but still mostly disordered, state. The compact FUS monomers can then interact with other monomers via exposed tyrosine and arginine residues to form droplets (Figure 6a). Importantly, the MS and MD data are in good agreement with recent observations from NMR that FUS adopts a compact state in droplets ^41^, which underscores the validity of our approach.

In case of TDP-43, both its folded NTD and the disordered C-terminal LCD have been implicated in LLPS ^44,45,53^. We find that phase separation of NT*-TDP-43 can be controlled by adjusting the pH, although this does not entail a clear shift in the distribution of charges along its sequence. There are virtually no titratable residues in the LCD of TDP-43, whereas the NTD requires a pH range from 10 – 5 to fold into its native structure ^54^. MS reveals the formation of dimers and trimers in a pH-dependent manner, in good agreement with the ability of the NTD to self-assemble, and supported by the presence of titratable residues at the domain interface ^44,54^. MS and MD simulations suggest that the protein, unlike FUS, remains at least partially flexible upon oligomerization, in line with the LCD being extended away from the folded pats of the protein. In the MD simulations, we observe a strong tendency for β-strand formation in the LCD, which again correlates well with the high fibrillation propensity of peptide fragments from this region ^55^. TDP-43 thus appears to adopt a segmented structure in droplets, where the NTDs form a scaffold from which the LCDs protrude to engage in low-specificity contacts within the same, or between different, oligomers (Figure 6a).

hCPEB3 exhibits yet another set of characteristics: nMS suggests that the protein remains fully flexible regardless of pH but is increasingly incorporated into fibrillar aggregates as the pH approaches the physiological range. Importantly, hCPEB3 has been suggested to undergo LLPS only in the presence of crowding agents ^16^, which may indicate a strong preference for ordered aggregation over disordered interactions. Its biological function as an engram of memory formation has early on been coupled to its assembly into stable structures which sequester mRNAs and thus produce a lasting impact on the cell’s translational profile ^56,57^. The high aggregation propensity indicates that the structure of hCPEB3 may be regulated by physical modifications such as SUMOylation, rather than weak interactions between disordered regions (Figure 6a).

nMS and MD not only inform about the respective assembly mechanisms of FUS, TDP-43, and hCPEB3, but also help to delineate the locations of their RRMs (Figure 6b). For FUS, the MD simulations do not suggest a specific orientation of the RRM in the compact state, and interactions between monomers would also be unlikely to point the RRMs into a specific direction. We therefore speculate that the orientations of FUS molecules in the droplet state would be dictated by the direction of bound RNAs, rather than the other way around. However, a different picture emerges for TDP-43. The “corkscrew” structure dictated by the N-terminal domains (Figure S4) would align the RRMs of neighboring protomers, which is also evident from our MD simulations. It was recently revealed that TDP-43 assembles into multimers along its target RNA sequences ^58^. In this scenario, NTD oligomerization could stabilize the protein-RNA complex in an extended state, while contacts between the LCDs promote their condensation into droplets, and potentially affect the accessibility of the bound RNAs. hCPEB3, on the other hand, forms fibrillar structures. The highly ordered nature of these assemblies implies that the RRMs are extended out from the fibril core, which was originally proposed by Kandel and co-workers for apCPEB. hCPEB3 may thus align its target RNAs and in this manner control their translation, a process known as vectorial channeling ^59^.

Our findings thus highlight the fact that hRNPs, despite being superficially similar RNA-binding proteins with disordered low-complexity domains, have evolved distinct assembly structures. By avoiding a one-size-fits-all mechanism, hRNPs can provide scaffolds with different properties, such the induction of different RNA conformations, in a highly specific manner.

## Materials and Methods

### Reagents

All chemicals were purchased from Sigma unless noted otherwise.

### Protein production

MiniMaSp1 was produced as described ^28^. Constructs containing a N-terminal NT* tag with 6-His repeat and a TEV cleavage site followed by either FUS, TDP-43 or hCPEB3 were ordered from Genscript. The plasmids were transformed into BL21(DE3) cells (Sigma) using heat-shock. Cells were grown in LB medium at 37°C and induced with IPTG at a OD_600_ at 0.6-0.8. After induction protein was expressed overnight at 18°C for NT*-FUS and NT*-TDP-43, and at 12°C for NT*-hCPEB3. Cells were harvested by centrifugation at 4000 rpm for 20 min at 4°C and stored at −80°C until purification. Prior to purification the cell pellet was thawed on ice and resuspended in 10 mL lysis buffer (dH_2_O with 10 mM imidazole, pH 9, 0.1 mg/mL lysozyme and cOmplete mini EDTA-free protease inhibitor tablets (Roche)) per 0.25 L culture before sonication at 50 % amplitude in 5 sec pulses for 10 minutes. Lysate was spun down at 10,000 g for 30 min at 4 °C, and the supernatant was loaded on a HisTrap column equilibrated in water with 10 mM imidazole, pH 9, washed with 1 M NaCl to reduce bound nucleic acids, and eluted the protein with imidazole (250 mM). Eluted fractions were analyzed on SDS-PAGE (miniProtean TGX, BioRad) under non-reducing conditions and without boiling. Fractions containing the protein of interest were dialyzed against dH_2_O prior to experiments, according to the ideal conditions identified for TDP-43 ^31^.

### Microscopy

Protein samples in dH_2_O were found to have a pH of 7.5, as measured by a microelectrode, which is due to residual histidine from the purification. By addition of 0.12 % or 12.5 % ammonia, the pH reached 10.5 and 12, as we previously used to study proteins under alkaline conditions ^39^. 50 μL of the solution were transferred to a nonbinding surface 96 well black plate with clear bottom (Corning). Samples were imaged at room temperature (ca. 23°C) on a CellObserver microscope (Zeiss) with the transmitted light channel. 20x or 40x magnification was used for imaging. For imaging of amyloids with the fluorophore Thioflavin T, we simply added Thioflavin T to the sample, and added a FITC excitation and emission filter set for imaging. Zen Blue (Zeiss) and ImageJ (https://imagej.nih.gov/) was used for analysis of the images.

### pH-dependent solubility assay

Concentrated NT*-tagged hRNPs were diluted in Tris buffers with pH 6, 7, 8, 9, 10 and 11 to reach a final buffer concentration of 20 mM tris and a final protein concentration of 1 mg/mL. Samples were incubated at room temperature, where aliquots were taken out at different time points, spun down at 13000 ×g for 10 min. Supernatant and pellet samples were mixed with 4x Laemmli sample buffer (non-reducing), and samples ran on a 4-15 % stain-free TGX gel (BioRad). The gels were imaged with ChemiDoc (BioRad).

### RNA binding assays

To assess whether NT*-FUS is co-purified with RNA, the final NT*-FUS samples (with or without a 1M NaCl wash during purification) were subjected to degradation of broadband RNase. 50 μL HisTrap elution fractions of NT*-FUS (1 μM protein, 255 mM imidazole) were first degraded using proteinase K (1 μg/μL final concentration) at 50°C for 45 minutes. After heat-denaturation of proteinase K (95°C, 15 minutes), 1 μL RNase A (30 units) was added to the NT*-FUS samples and incubated at 37°C for 30 minutes. Yeast tRNA extract (Sigma, 10 μM in 50 mM imidazole) served as a control sample. The reactions were analyzed on denaturing polyacrylamide gel electrophoresis (Mini-PROTEAN TBA-Urea Precast Gels, BioRad), run for 30 minutes at 250 V. 5 μL sample was diluted in 5 μL formamide containing bromphenol blue, heated to 95°C for 2 minutes and loaded onto the gel. The gel was stained with SYBRGold (Sigma) according to manufacturer instructions and imaged with ImageQuant 800 (Amersham).

### Native and ion mobility mass spectrometry

NT*-tagged hRNPs in dH_2_O containing 0, 0.125 or 12.5 % ammonia to reach pH 7.5, 10.5 or 12.0 with a final protein concentration of 10 to 15 μM were mixed immediately prior to transfer to a nESI capillary (Thermo). Mass spectra were acquired in positive ionization mode on a Waters Synapt G1 TWIMS MS modified for analysis of intact protein complexes (MS Vision, The Netherlands), equipped with an off-line nanospray source. The capillary voltage was set to 1.5 kV. The trap voltage was set to 10 V and stepwise increased up to 100 V for collisional activation. The source temperature was 30 °C. The cone voltage was 50 V, and the source pressure was set to 8 mbar. The ion trap voltage was set to 10 V and increased in steps of 10 V to 150 V for collision induced unfolding. Trap gas was argon with a flow rate of 4 mL/h, and IMS gas was nitrogen with a flow rate of 30 mL/h. IMS settings were wave height 10 V, wave velocity 200 V for NT*-TDP-43, and wave height 11 V, wave velocity 450 V for NT*-FUS. β-lactoglobulin, BSA, and Concanavalin A (Sigma) were used as T-Wave calibrants for NT*-TDP43, and BSA only (due to the virtually identical MW) for NT*-FUS as described ^60^. Data were analyzed using MassLynx 4.0 and Pulsar software packages ^61^.

### CCS database searches

To find candidate shapes for the TDP-43 dimer, PDB searches were performed as described ^42,62^, using a CCS of 6990 Å^2^ and a MW of 122 kDa as search criteria. The ten best-scoring homodimers were extracted from the list of the 50 best matches, and the aspect ratios were calculated in UCSF Chimera ^42,63^.

### Modelling and simulation of NT*-hCPEB3^1-40^

The solution state structure of the N-terminal domain (NT) from the spider silk protein was retrieved from the Protein Data Bank (PDB ID: 2LPJ). The lowest energy state structure (Model 1) from the ensemble of NT structures was selected and two *in-silico* mutations “D40K and K65D” were incorporated (NT*) into the structure. A 47 amino acid sequence comprising of TEV cleavage site (^1^ENLYFQS^7^) and the first 40 residues from human CPEB3 protein (Uniprot ID: Q8NE35) was considered for modelling. Extended state conformation of TEV+CPEB3^1-40^ polypeptide segment was generated using the LEaP module of AMBER18^64^, and attached via a peptide bond with NT*. The NT*-TEV-CPEB3^1-40^ construct was then subjected to explicit solvent MD simulations. The N- and C-terminus of the chimeric model were capped with ACE (acetyl) and NME (N-methyl) functional groups respectively. The model was placed in the centre of a truncated octahedral box whose dimensions was fixed by maintaining a minimum distance of 8 Å between any protein atom and the box boundaries. TIP3P water model ^65^ was used to solvate the structure. The net charge of the systems was neutralized by adding 8 Na^+^ counter ions. All-atom molecular dynamics simulations were carried out using the PMEMD module through the AMBER18 suite of programs employing ff14SB force field parameters ^66^. The system was energy minimized using steepest descent followed by conjugate gradient schemes, heated to 300 K over 30 ps under NVT ensemble and equilibrated for 200 ps in NPT ensemble. The final production dynamics was run for 1 μs under NPT conditions. Simulation temperature was maintained at 300 K using Langevin dynamics ^67^ (collision frequency: 1.0 ps^−1^) and the pressure was kept at 1 atm using weak-coupling ^68^ (relaxation time: 1ps). Periodic boundary conditions were applied on the box edges and Particle Mesh Ewald (PME) method ^69^ was used to compute long range electrostatic interactions. SHAKE algorithm ^70^ was used to constrain all the bonds involving hydrogen atoms and the equation of motion was solved with an integration time step of 2 fs.

### Modelling and simulation of FUS

The initial full length unrelaxed structure of FUS (526 residues; Uniprot ID: P35637) was generated using Alphafold2 (AF2) through the jupyter notebook for Colabfold ^71^ using the default parameters and no template. The AF2 generated protein model was then placed in the centre of a truncated octahedral box whose dimensions was fixed by maintaining a minimum distance of 6 Å between any protein atom and the box boundaries. OPC ^72^, which is a 4-point rigid water model was used to solvate the structure and the net charge of the systems was neutralized by adding 14 Cl^-^ counter ions. The final system consisted of a total of 251 160 atoms. All-atom molecular dynamics simulations were carried out using the PMEMD module through the AMBER18 suite of programs employing ff19SB force field parameters ^73^. The system was energy minimized using steepest descent followed by conjugate gradient schemes, heated to 300 K over 30 ps under NVT ensemble and equilibrated for 200 ps in NPT ensemble. The final production dynamics was run for 500 ns under NPT conditions.

### Modelling and simulation of TDP-43

The initial full length structure of TDP-43 monomer was the SAXS-derived MD model (418 residues), provided by Prof. Samar Hasnain, Liverpool. AF2 was used to generate oligomeric structures of the TDP-43 N-terminal domains (NTD). Then the full-length monomer model was fitted into the AF2 structures of NTD oligomers to generate full length TDP-43 dimers and trimers. Coarse-grained molecular dynamics simulations for TDP-43 monomer, dimer or trimer were carried out using the AWSEM force field ^74^, running in openMM with Langevin integrator with a time step of 2 fs. AWSEM annealing was iterated twice to achieve a consistent prediction. In each annealing generation, simulations started with the same initial structures and were run 20 times with different random seeds. For the 1st generation, systems cooled down from 400K to 300K in 6×10^6^ steps (around 12 μs in laboratory time). Then, the last frames of annealing were relaxed for another 6×10^6^ steps at 300K. For the 2nd generation, those contacts that were formed more than 6 times in the 20 structures from the last frames of the relaxing simulations in the 1st generation, were then used as constraints during annealing. Here, a contact refers to a residue pair with their CA atoms distance smaller than 9.5 A. The annealing and relaxing protocol was the same as for the 1st generation runs. Finally, the last frames of the 2nd generation runs became the end structures. Domain contact maps (Fig. S4b) were calculated by counting the average number of contacted residue pairs between TDP-43 domains over 20 end structures. The average and standard variance of electrostatic energies per protomer for 20 end structures (Fig. S4d) were calculated using the definition of the Debye-Huckel potential in the AWSEM Hamiltonian, under different pH conditions.

### Negative stain electron microscopy

For negative stain EM, 4 mg of lyophilized hCPEB3 were dissolved in ddH_2_O, sonicated for 30 sec and incubated for 3 days at 37 °C. The CPEB3 samples were either diluted in water or in 10 mM HEPES pH 7.15, 75 mM NaCl, 10 mM KCl, 2 mM MgCl2 and 5 mM TCEP ^48^ to a final concentration of 2 to 4 μM CPEB3. 5 μL was loaded to 200 mesh copper grids with Formvar/Carbon support film that had been glow-discharged at 25 mA for 30 sec. After one minute of incubation, the liquid was removed and the negative staining was carried out by applying a 5 μl of 1 % (w/v) uranyl acetate (in H_2_O) to the grid for 20-30 seconds. Then, the liquid was removed and the procedure was repeated 6 times. The grids were imaged in Talos 120 C G2 (Thermo Scientific) equipped with a CETA-D detector.

## Acknowledgements

CS is supported by a Novo Nordisk Foundation Postdoctoral Fellowship (NNF19OC0055700). DL is supported by a Swedish Research Council grant for Internationally Recruited Scientists (2013-08807) to Prof. Sir David P. Lane, Karolinska Institutet. EGM is supported by a project grant from the Swedish Research Council (2020-04825). AR is supported by the European Research Council (ERC) under the European Union’s Horizon 2020 research and innovation program (grant agreement No 815357), the Swedish Research Council (2019-01257) and Formas (2019-00427). The contributions from XG and WGP were supported by the Center for Theoretical Biological Physics, sponsored by NSF grants CHE-1743392 and PHY-2019745. Additionally, we wish to recognize the D.R. Bullard Welch Chair at Rice University, Grant C-0016 (to PGW). ML is supported by a KI faculty-funded Career Position, a KI-Cancer Blue Sky grant, an Ingvar Carlsson Award from the Foundation for Strategic Research (SSF), a Cancerfonden Project grant (19 0480), and a VR Starting Grant (2019-01961). We gratefully acknowledge support from Dr. Florian Salomons and the Biomedicum Imaging Core (BIC) at Karolinska Institutet. Special thanks are extended to Prof. Nancy M. Bonini (University of Pennsylvania) and Prof. Justin Benesch and Prof. Andrew J. Baldwin (University of Oxford) for helpful discussions.

## Data availability statement

All data needed to evaluate the conclusions in the paper are present in the paper and/or the Supplementary Materials.

