## Supplemental Information for "Mass spectrometry of RNA-binding proteins during liquid-liquid phase separation reveals distinct assembly mechanisms and droplet architectures"

**This PDF file includes:**

Supplementary Text
Figs. S1 to S6
Movie S1
References (75 to 79)

### Supplementary Text

Effects of the NT*-domain on hRNP conformation and self-assembly

In this study, we find that NT* enables the purification of hRNPs under non-denaturing conditions. Previous work from us and others has established that the NT* domain is an efficient solubility tag [30,75,76]. Intracellular hydrogen/deuterium exchange studies have revealed that NT* remains folded even if fused to aggregation-prone proteins, thus preventing the formation of insoluble aggregates [77]. The proteins included here display different aggregation propensities *in vitro*. TDP-43, for example, aggregates upon tag removal [21], whereas hCPEB3 is soluble only under denaturing conditions [78]. Fusion to the NT* domain enables us to purify and study all three proteins under identical solution and nMS conditions, but also raises the question to what extent it affects their structures and self-assembly. In TDP-43, the NT* domain is connected to a folded NTD via a short, disordered segment composed of the TEV cleavage site and the first 4-5 residues of TDP-43. We detect by nMS dimerization and trimerization of NT*-TDP-43 at pH 7.5, as predicted for isolated TDP-43 NTDs. NT*, on the other hand, does not oligomerize [30]. We thus conclude that our observations can be attributed to specific oligomerization of TDP-43, although we cannot exclude that the NT* domain may influence the efficiency of the process. Recently, we found that NT* induces compaction of the first 60 residues of the disordered transactivation domain of p53 by binding to short hydrophobic segments [79]. As for p53, NT* is in hCPEB3 and FUS connected to a disordered N-terminal domain; however, the disordered NTD of FUS is three times longer than that of p53 (270 *vs.* 90 residues) and has a significantly lower hydrophobicity score (Figure S6). We thus consider it highly unlikely that the near-complete unfolded-to-globular transition we observe in FUS can be attributed to the presence of the NT* domain. For hCPEB3, we do not observe any compaction comparable to that of p53 or FUS, which is in agreement with the very transient contacts observed in MD simulations (Figure S6). Based on this evidence, we conclude that the NT* domain reduces the aggregation propensity of hRNPs through steric hindrance and its own high solubility but does not prevent low-complexity domains from forming transient interactions that mediate LLPS. As a result, the different structural plasticities observed here are specific for each protein.

.

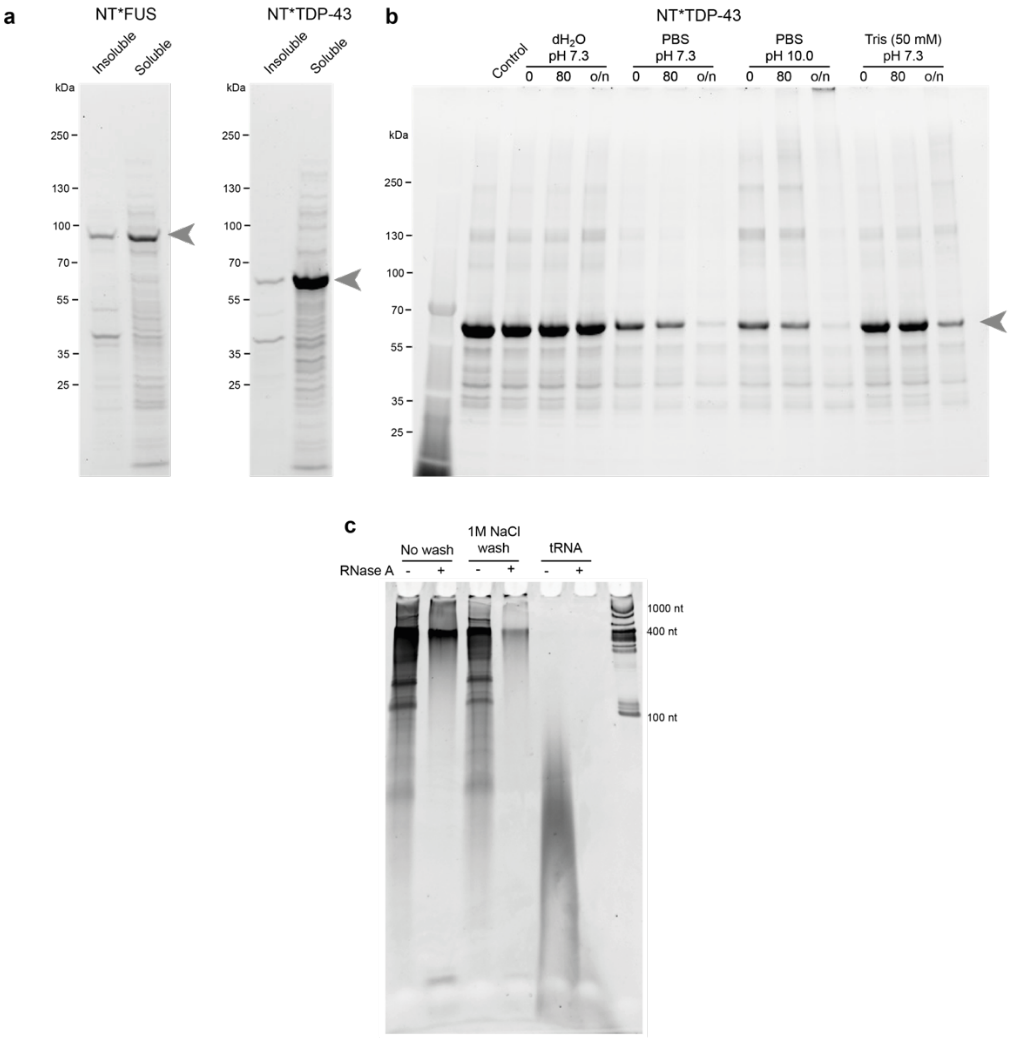


**Figure S1. Purification and characterization of NT*-tagged FUS and TDP-43.** (a) NT*-FUS and NT*-TDP-43 expressed in *E. coli* BL21 under standard conditions (see Methods) is located predominantly in the soluble fraction following lysis by gentle sonication in $dH_2O$. (b) Stability of NT*-TDP-43 in standard buffer shows moderate aggregation after prolonged incubation in PBS and Tris, and good stability in $dH_2O$. (c) Denaturing PAGE of NT*-FUS purified by affinity chromatography under native conditions reveals co-purified endogenous RNAs.

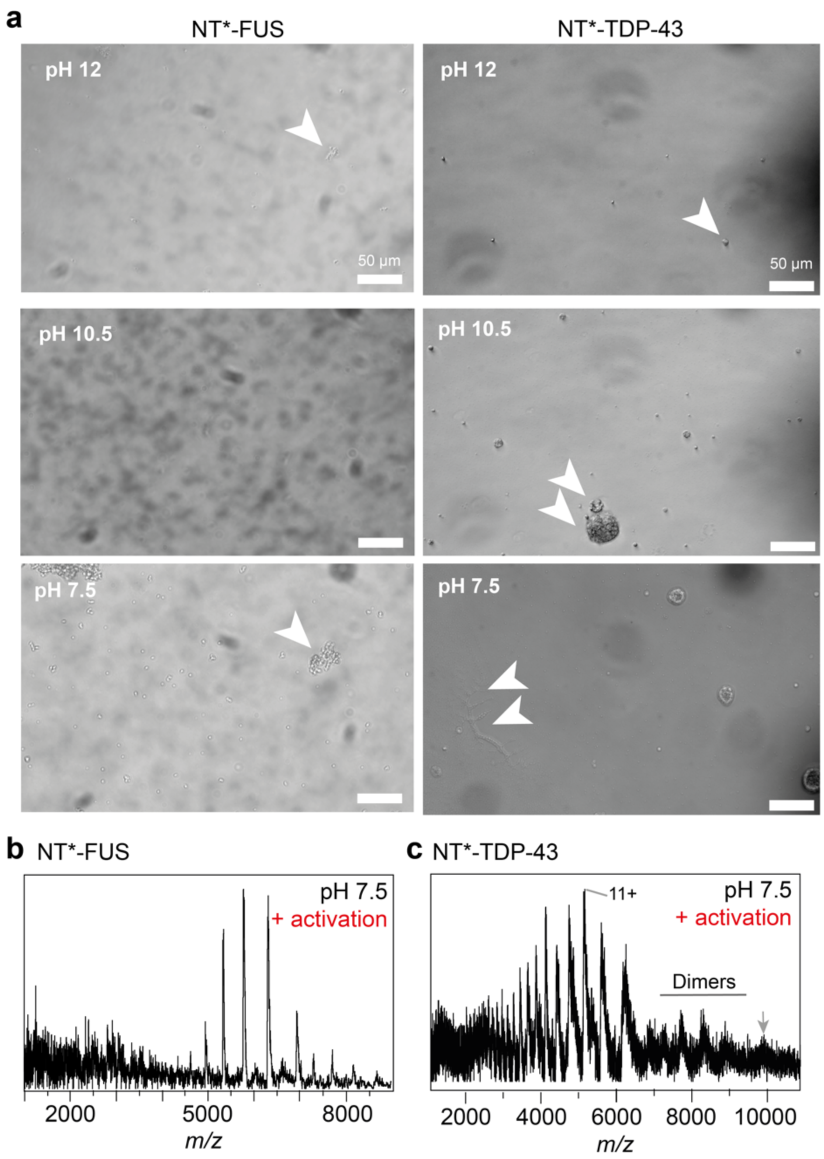


**Figure S2. Bright-field microscopy and nMS of NT*-FUS and NT*-TDP-43 assemblies.** (a) Brightfield microscopy images of NT*-FUS and NT*-TDP-43 at pH 12, 10.5, and 7.5. We observe small amounts of amorphous aggregates (arrows) at all pH values, which likely represent protein carried over from the insoluble fraction during purification. (b) and (c) nMS spectra of NT*-FUS and NT*-TDP-43, respectively, at pH 8 with an ion trap energy of 100 V. Collisional activation releases monomeric NT*-FUS with a lower charge than free protein (compare Figure 2b). Collisional activation of NT*-TDP-43 at pH 7.5 does not have a notable impact on protein charge states compared to pH 10.5 (compare Figure 2c). The arrow indicates the putative 19+ charge state of the trimer.

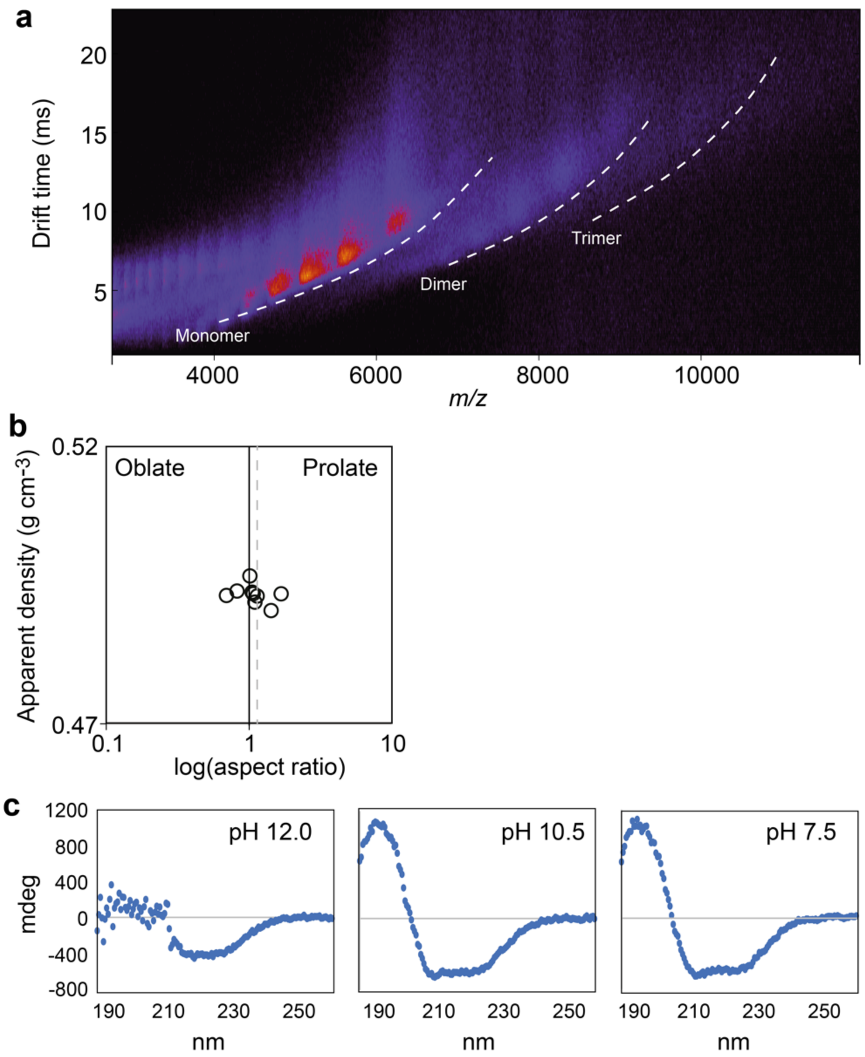


**Figure S3. Characterization of NT*-TDP-43.** (a) The mobiligram of NT*-TDP-43 at pH 7.5 shows ion series corresponding in mass to compact monomers, dimers, and trimers. (b) Searching the PDB for protein dimers with similar MW and CCS as dimeric NT*-TDP-43 yields both oblate and prolate structures with an aspect ratio close to 1, suggesting a mixture of possible shapes for the dimer. The average aspect ratio for the top ten matches is indicated by a dashed line. (c) CD measurements of NT*-TDP-43 at alkaline and physiological pH show folding at pH 10 and below but no pronounced changes between pH 10 (onset of droplet formation) and pH 7.5 (complete droplet formation), which indicates that LLPS occurs after the protein has folded.

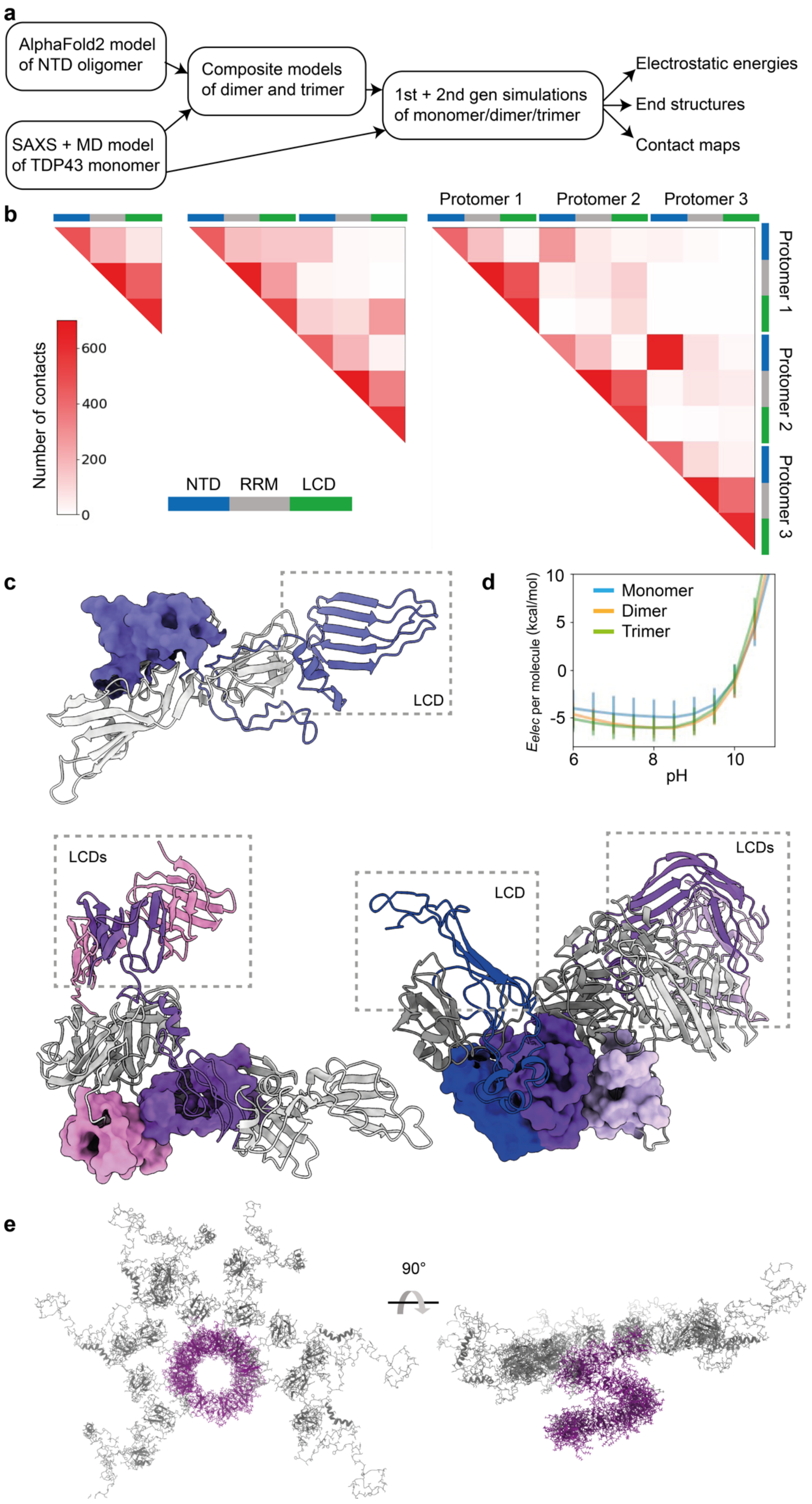

a
AlphaFold2 model of NTD oligomer
SAXS + MD model of TDP43 monomer
Composite models of dimer and trimer
1st + 2nd gen simulations of monomer/dimer/trimer
Electrostatic energies
End structures
Contact maps
b
Protomer 1
Protomer 2
Protomer 3
Number of contacts
600
400
200
0
NTD
RRM
LCD
c
LCD
LCDs
LCD
LCDs
d
Monomer
Dimer
Trimer
$E_{elec}$ per molecule (kcal/mol)
pH
e
90°

**Figure S4. Simulations of TDP-43 oligomers.** (a) The computational strategy employed to generate TDP-43 monomer, dimer, and trimer structures. Briefly, AF2 was used to generate oligomeric structures of the TDP-43 NTD. SAXS-derived MD models (kindly provided by Prof. Samar Hasnain, Liverpool) were then fitted into the NTD oligomers to generate dimers and trimers. Monomers, dimers, and trimers were subjected to two rounds of cooling and relaxing in 20 replicates each (see methods) to obtain end structures for electrostatic energy and contact map calculations. (b) Contact maps of monomers, dimers and trimers show extensive contacts between the NTDs and RRMs, and RRMs and LCDs, but not the NTDs and LCDs. (c) Representative example structures of monomers, dimers, and trimers. The NTDs are rendered as surfaces, the RRMs as grey cartoons, and the LCDs as colored cartoons. (d) Electrostatic energy calculations of the monomers, dimers, and trimers at alkaline and neutral pH reveal slightly more favorable energies in the dimers and trimers which is likely due to NTD contacts absent in the monomer. (e) View of the complete TDP-43 oligomer model. The first 8 TDP-43 molecules are shown as full-length, the full NTD "corkscrew" (purple) is shown for 14 molecules.

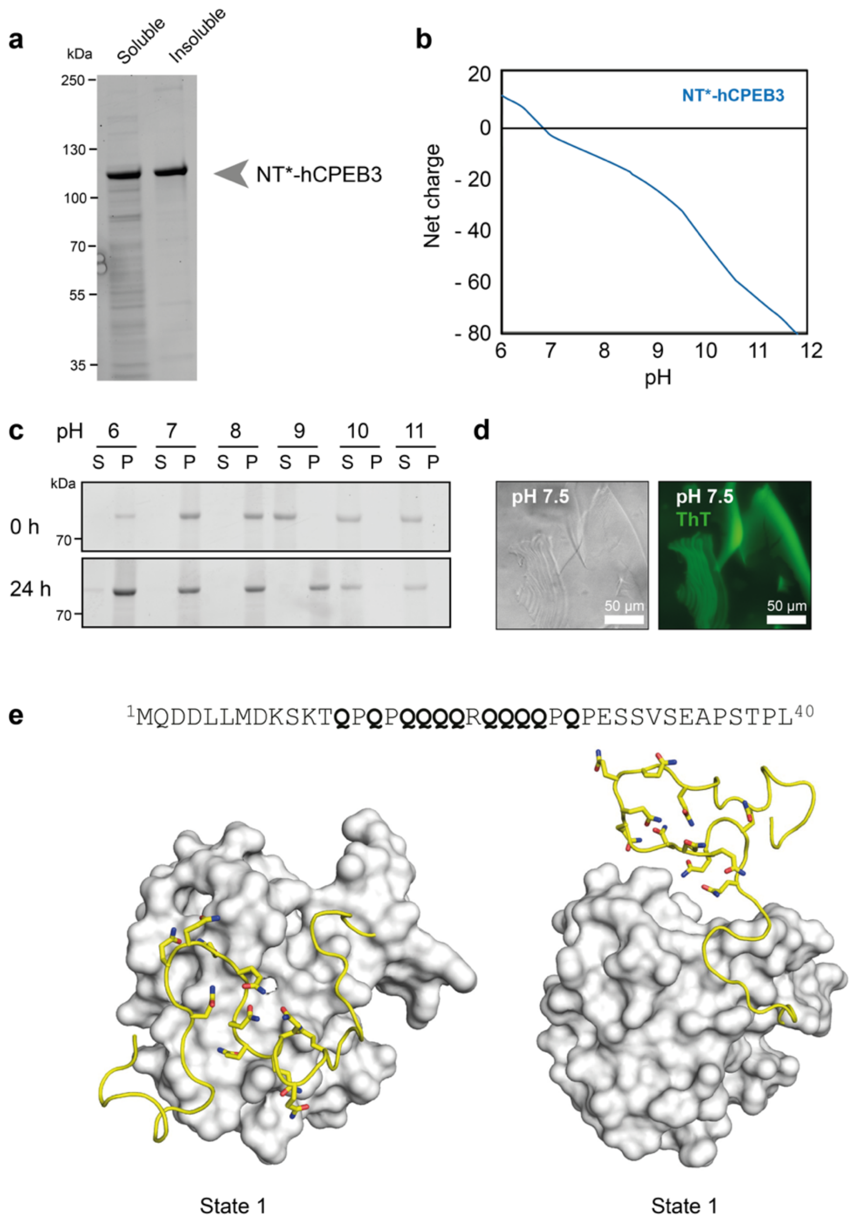


**Figure S5. Characterization and interactions in NT*-hCPEB3.** (a) Expression of NT*-tagged hCPEB3 yields partially soluble protein as judged by SDS-PAGE. (b) pH-dependent solubility assay shows insoluble NT*-hCPEB3 below pH 8. After 24h, the protein also became insoluble at pH 9. (c) Computed net charge of NT*-hCPEB3 from pH 6 to 12. The experimentally observed pH for aggregation is indicated by a shaded area. (d) Fluorescence microscopy of the hCPEB3 assemblies visible by light microscopy shows staining for the amyloid-specific dye ThT, confirming the presence of fibrillar material. (e) MD simulations of NT* fused to the first 40 residues of hCPEB3. The hCPEB3 sequence is shown above, with the polyglutamine sequence highlighted. Two representative snapshots from the simulations show the polyglutamine stretch bound to the surface of NT* (state 1) or in a flexible conformation facing away from NT* (state 2). The surface of NT* is rendered in grey, the glutamine residues are shown as sticks.

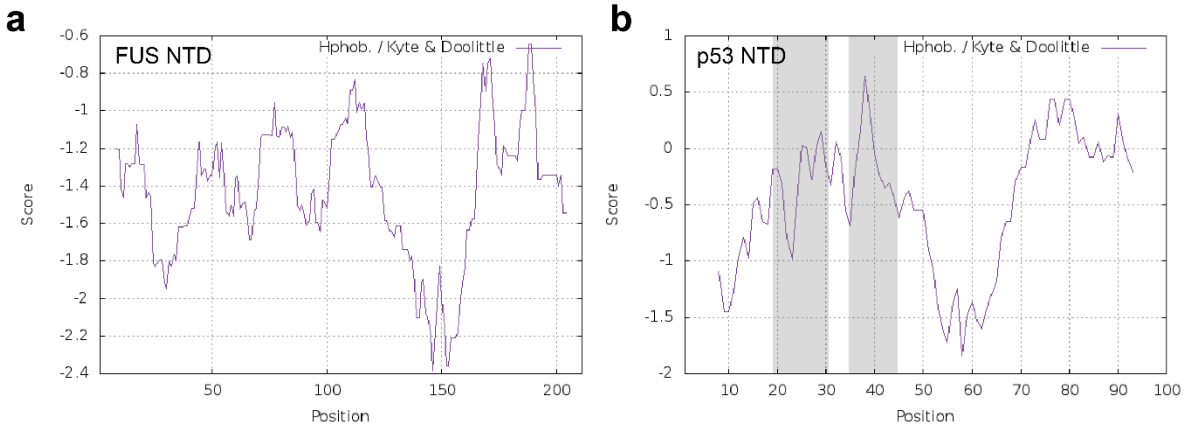


**Figure S6. Hydrophobicity profiles of the NTDs from FUS and p53.** Average hydrophobicities with a window size of 9 residues were computed using the ProtScale server (https://web.expasy.org/protscale/). The p53 segments that bind to the surface of NT* are indicated by grey boxes.

**Movie S1.**

NT*-FUS diluted from a pH 9 dH2O stock into 20 mM Tris pH 6, 500 mM NaCl, shows phase separation into spherical droplets that fuse on a minute timescale (total 15 minutes). Imaged every 5 sec for 181 frames. Scale bar is 50 µm.
